# FUSION PORE CLOSURE DETERMINES SUBQUANTAL KISS-AND-RUN SEROTONIN RELEASE FROM FULLY LOADED DENSE-CORE VESICLES

**DOI:** 10.64898/2026.11.05.764715

**Authors:** Francisco F. De-Miguel

## Abstract

During kiss-and-run exocytosis, a transient fusion pore opens, releases part of the vesicular quantum, and closes without full fusion. This report investigates two questions: first, whether dense-core vesicles undergoing kiss-and-run contain the same number of molecules as those undergoing full fusion; second, whether subquantal release results from pore closure or vesicle depletion. We analyzed amperometric recordings of extrasynaptic serotonin release from dense-core vesicles in cultured leech Retzius neurons. Full-fusion events released 91,585 ± 34,428 molecules (*n* = 14), whereas kiss- and-run events released 58,344 ± 20,759 molecules— 64% of the quantum. We analyzed the biexponential dynamics of the filtered amperometric current using Ohmic equations for cylindrical and toroidal pores. Hydrated flux packing (HFP) independently estimated the minimum peak pore diameter from the number of hydrated serotonin molecules crossing the pore. Kiss-and-run peak diameters were 2.1–3.2 nm, corresponding to the passage of 4–10 hydrated molecules. The vesicular serotonin content of individual kiss-and-run events was estimated by a frozen pore extrapolation (FPE), in which pores were kept at the peak HFP diameter. FPE yielded 102,650 ± 10,989 molecules per vesicle (*n* = 15), similar to the quantum obtained by integrating full-fusion events. Therefore, kiss-and-run subquantal release did not reflect a smaller quantum. We then compared the decay of serotonin flux with that estimated through the frozen pore at its peak diameter. In every event, the flux decayed faster, reaching half its peak in 0.59 (0.23–0.95) of the time taken by the frozen pore flux. We conclude that subquantal release during kiss-and-run exocytosis is governed by fusion-pore closure rather than vesicle depletion.

**Summary:** Estimates of individual vesicle quantum, the dynamics of serotonin release and pore closure confirm that subquantal release of serotonin during kiss-and-run exocytosis is due to closure of the fusion pore and not to transmitter depletion.

## Introduction

A nanometer-sized fusion pore opens within a fraction of a millisecond and may then expand (Heuser et al., 1979; Chandler and Heuser, 1980; Breckenridge and Almers, 1987a; Zimmerberg et al., 1987). The nascent pore allows water into the vesicle, hydrates the intravesicular matrix and lets soluble transmitter leak into the extracellular space (Chow et al., 1992; Wightman et al., 1995; Staal et al., 2004; Amatore et al., 2005). Usually the pore dilates and the vesicle fuses fully, releasing the rest of its quantum. In other cases the pore closes again (Breckenridge and Almers, 1987b; Wightman et al., 1991; Alvarez de Toledo et al., 1993). This mode, known as kiss-and-run (Ceccarelli et al., 1973; Alabi and Tsien, 2013), releases only a fraction of the quantum (MacDonald et al., 2006; Omiatek et al., 2010).

Subquantal release could arise in two ways. The pore could close before the vesicle has emptied, or it could remain open while release slows down because the vesicle is depleted and the concentration gradient across the pore falls. The two mechanisms predict different time courses of the amperometric current after its peak. Closing pores produce faster decay of release. Distinguishing them requires determining the size of the pore and the transmitter content of the vesicle, neither of which is measured directly by amperometry.

Retzius neurons of the leech release, from their somata quanta of about 90,000 serotonin molecules from large dense-core vesicles (Trueta et al., 2003; Bruns et al., 2000). Here we analyzed amperometric kiss-and-run events from extrasynaptic dense-core vesicles in these neurons in three steps. First, we introduced a method to estimate the minimum pore diameter from the serotonin flux, based on the optimal hydrated flux packing (HFP) of molecules in a circular section. Second, to determine if the decay of serotonin flux indicates pore closure or depletion, we estimated the decay of the serotonin content through pores frozen at their peak. Third, to confirm that subquantal release is produced by pore closure, we estimated individual quantal serotonin content using a frozen-pore extrapolation (FPE). For that, the pore was kept at the peak HFP diameter through an equation that linked it to its effective length and vesicle volume. The vesicular content measured this way was compared to that of full-fusion events. We found that pores undergoing kiss-and-run close faster than the vesicle empties.

## Materials and methods

### Isolation and culture of neurons

Experiments were conducted on serotonergic Retzius neurons isolated from leeches (*Hirudo verbana*). Individual neurons were removed from the ganglion one by one with a glass pipette, then sterilized and plated in culture (Dietzel et al., 1986; Trueta et al., 2003). Experiments were performed at 18°C after 1–8 days in culture.

### Amperometric recordings of serotonin release

Amperometric currents produced by serotonin oxidation were recorded with a carbon fiber electrode of 10 µm tip diameter held at 650 mV. The tip of the carbon fiber was positioned touching gently the primary axon of the neuron under an inverted microscope (Nikon Eclipse TE200). Exocytosis was evoked by mechanical pressure applied to the membrane through the carbon fiber. Currents were recorded with an Axopatch 200B amplifier (Molecular Devices), filtered at 10 kHz (low-pass Bessel), digitized at 100 kHz and stored with pClamp 9 (Molecular Devices). The membrane potential of the neuron was recorded with an intracellular microelectrode connected to an Axoclamp 2B amplifier after compensation for the electrode resistance.

### Spatial resolution of the recordings

The kinetics of amperometric currents are distorted when transmitter must diffuse a long distance before reaching the electrode. The electrode tip was therefore placed in contact with the membrane, and the pressure used to stimulate exocytosis probably reduced the distance between release sites and the carbon surface further. Even so, diffusion between the pore and the electrode smooths the current over a fraction of a millisecond (Wightman et al., 1995; Zhang and Jackson, 2008), so that the earliest phase of the current, which also reflects hydration of the intravesicular matrix (Alvarez de Toledo et al., 1993; Hafez et al., 2005), was not used to infer pore dynamics.

### Event selection and filtering

Amperometric spikes were detected with Quanta Analysis 8.2 (Sulzer laboratory, Columbia University) running in Igor Pro 6.0 (WaveMetrics); transients exceeding the standard deviation of the baseline noise by more than fivefold and free of overlap with other spikes were included. Selected currents, either full-fusion spikes (≥10 pA) or small kiss-and-run events (≤6 pA), were filtered with statistical empirical mode decomposition (SEMD), which includes a smoothing step (Kim et al., 2012). Kiss-and-run events were defined as small spikelets with fast rise and an exponential decay (see Results). Fifteen kiss-and-run events and 14 full-fusion events were analysed here.

### Empirical description of the currents

Each current was fitted to:

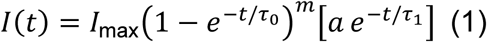

where *I*_max_ is a scale factor, the first factor describes the rise with a time constant *τ*_0_ and exponent *m*, and the bracketed term describes the decay. The fitted currents were used for all subsequent calculations and extrapolated with exponential tails to 100 ms.

### Serotonin flux and cumulative release

By Faraday’s law, the serotonin flux ***J***(***t***) and the number of molecules released up to time *t*, *q*(*t*), are

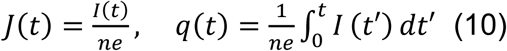

where *n* = 4 is the number of electrons released per oxidation of one serotonin molecule (Bruns et al., 2000) and *e* is the elementary charge. Because serotonin is oxidized at the electrode, its extravesicular concentration was taken as zero. From Fig. 3 onward, amperometric currents are presented as serotonin flux, *J* = *I*/*ne*, in molecules per microsecond, so that *I*_peak_τ/*ne* in Eq. 12 equals *J*_peak_τ.

### Pore diameter from hydrated flux packing (HFP)

To estimate the minimum pore diameter compatible with the measured flux, we determined the minimum circular cross-section required to accommodate the number of hydrated serotonin molecules crossing it per microsecond. The flux at the current peak was expressed in molecules per microsecond, *N* = *J*(*t*_peak_) × 10^−6^ s, and rounded up to the next integer, as a fractional value implies that more molecules than the lower integer must pass.

Each serotonin molecule crossing the pore was represented as a sphere of hydrated diameter *d*_H_ = 0.8–0.9 nm (central value 0.85 nm). Four lines of evidence support the hydrated rather than the bare diameter. First, serotonin crosses the pore as a cation. Its apparent p*K*a is 10.4 ± 0.2 (Chattopadhyay et al., 1996), so more than 99.9% of the molecules carry a protonated ammonium group both in the acidic vesicle lumen and at extracellular pH. This charged group binds hydration water strongly. Second, the size of a hydrated monoamine cation follows from its diffusion coefficient in free solution. The Stokes–Einstein diameters, *d* = *k*_B_*T*/(3π*ηD*), of the five biogenic amine cations measured by Rice et al. (1985) are 0.88–0.97 nm. The molecular masses of these five amines (153–183 Da) bracket that of serotonin (176 Da). The range used here lies at or below the lower end of those values, so that the HFP diameters remain lower bounds. Third, serotonin consists of a planar 5-hydroxyindole ring with a gauche–gauche ethylamine side chain (Naeem et al., 2022). The orientation of serotonin is expected to be randomized within tens of picoseconds, as that of the indole amino acid tryptophan in water (46 ps; Lu et al., 2004), far faster than the 1-µs window over which HFP counts molecules. Its effective cross-section is therefore the rotationally averaged size of the hydrated molecule (the Stokes–Einstein diameter), rather than the 0.67-nm sphere of equal van der Waals volume (155 Å³; Zhao et al., 2003). Fourth, permeation involves dehydration only in channels whose lumen approaches the size of the bare solute and whose walls provide coordinating groups that replace the lost water (Hille, 2001); a lipidic fusion pore 2–3 nm wide, three to four times the hydrated diameter, meets neither condition.

For comparison, the bare diameter of serotonin was taken as that of the sphere of equal van der Waals volume (0.67 nm; see above). The minimum enclosing diameter of *N* non-overlapping circles was obtained from the circle packing in a circle problem (Graham et al., 1998): for *N* ≤ 9, the optimal arrangements are rings, with or without a central circle, possessing closed-form solutions; for *N* = 10, the arrangement was determined by constrained numerical optimization (*R*/*r* = 3.813, the proven optimum; Pirl, 1969). The HFP diameter is given by:

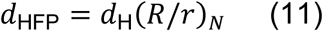

where (*R*/*r*)*_N_* is the ratio between the radius of the enclosing circle and that of each molecule in the optimal packing of *N* circles: 1, 2, 2.15, 2.4, 2.7, 3, 3, 3.3, 3.6 and 3.8 for *N* = 1–10 (Graham et al., 1998). Because seven circles fit into the same enclosing circle as six, *N* = 6 and *N* = 7 yield the same diameter. For each *N*, Eq. 11 defines the tightest arrangement (the minimum diameter); the maximum diameter that still corresponds to ***N*** is the threshold at which *N* + 1 molecules can fit, *d*_H_(*R*/*r*)*_N_*_+1_. The range of hydrated diameters (0.8–0.9 nm) introduces an uncertainty to both values. HFP relies solely on the measured current and the hydrated molecular size, requiring neither a diffusion coefficient nor the vesicular content.

### Vesicular content estimated by frozen pore extrapolation (FPE)

The quantal vesicular content *N*_quant_was measured as the time integral of full-fusion spikes (Eq. 10 for *t* → ∞). For kiss-and-run events, in which release is incomplete, the total content was estimated by assuming that, from the current peak onward, the pore remains frozen at its HFP diameter, so that the efflux of serotonin is set by the effective length of the pore, *L*_eff_, its area and the vesicle volume (Eq. 13). This procedure, frozen pore extrapolation (FPE), follows the equation:

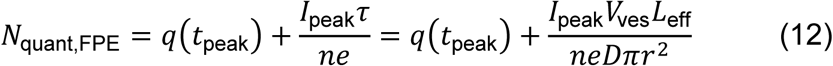

where ***τ*** is the emptying time constant of the frozen pore (Eq. 13) and *r* = *d*_HFP_/2. This expression represents the condition under which a fixed pore reproduces the measured peak current. The FPE contents (102,650 ± 10,989 molecules; range 88,110–121,707) were used in all analyses that require the vesicular content, including the reconstruction of pore diameters (Fig. 4) and the separation of closure and depletion (Eq. 14). Because the amperometric current fixes only the product of *D* and the vesicular content, the FPE contents are conditional on *D*_common_, which was calibrated in the companion paper on the HFP diameter of the median event; an error in *D* would scale all contents by the same factor (Supplemental text, S6). Their scale was therefore checked independently against the mean quantum measured from full-fusion events (91,585 molecules; Results).

### Release through a pore fixed at its peak diameter

If the pore remained frozen at its peak diameter *d*, release would follow first-order kinetics. For a well-mixed vesicle of volume *V*_ves_and a perfect extracellular sink, Fick’s law gives:

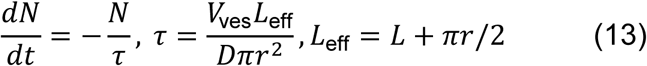

where *N*is the number of molecules remaining in the vesicle, *r* = *d*/2, *L*_eff_ is the effective length of a cylindrical pore of length *L*, including the access resistance of its two mouths (Hall, 1975; Hille, 2001), and *V*_ves_ = 5.84 × 10^−16^ cm^3^, the volume of a vesicle 103.7 nmin diameter (the mean corrected diameter of dense-core vesicles in Retzius neurons; Bruns et al., 2000). We used the diffusion coefficient common to toroidal and cylindrical reconstructions of these events, *D*_common_ = 4.49 × 10^−7^ cm^2^/s (see companion paper), with its accompanying cylinder length, *L* = 4.1 nm (companion paper); a toroidal pore with a rim curvature radius of **7**. **37** nmand the same coefficient gives time constants within 2.4%. The fixed-pore current from the peak onward is then *I*_fix_(*t*) = *I*_peak_ exp[−(*t* − *t*_peak_)/*τ*]. Note that ***τ*** depends solely on the pore geometry, the vesicle volume and *D*, not on the vesicular content (Supplemental text, S2). For each event, *τ* was evaluated at *d* = *d*_HFP_, and the recorded and fixed-pore currents were compared through their times to decay to one-half and one-quarter of the peak value.

### Contributions of closure and depletion to the decay

Because *I*(*t*) = *ne* [*N*_quant_ − *q*(*t*)]/[*V*_ves_*R*(*t*)], where *R*(*t*) = *L*_eff_/(*D*π*r*^2^) is the diffusional resistance of the pore, the fall of the current after the peak can be separated into a depletion term and a pore-resistance term:

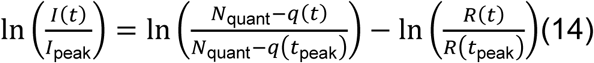

The fraction of the current decrease at half-decay attributable to the increase in *R* was defined as the closure share.

### Dynamics of toroidal and cylindrical pores

Equations describing the dynamics of toroidal and cylindrical pores are described in the companion paper.

### Statistical analysis

Values are expressed as mean ± SD with ranges. Kiss-and-run and full-fusion releases were compared using Welch’s *t*-test and the Mann–Whitney *U* test; normality was assessed with the Shapiro–Wilk test. Paired comparisons between recorded and fixed-pore decay times used the two-sided exact Wilcoxon signed-rank test, and associations were evaluated using Spearman’s rank correlation. Exact *p*-values are reported throughout. Calculations were performed in Python 3.11 using NumPy and SciPy.

## Results

### Kiss-and-run events release a fraction of the quantum

Gentle pressure applied to the primary axon with the carbon fiber evoked bursts of amperometric spikes. Most were large spikes whose amplitude (11.07 ± 2.62 pA) and fast rise time (0.95 ± 0.15 ms) resembled those of full fusion of dense-core vesicles in these neurons (Bruns et al., 2000). Thirteen percent of the events were smaller spikelets that rose rapidly and decayed along a single exponential, appearing as a straight line on semilogarithmic axes (Fig. 1 A). Fifteen spikelets analyzed had peak currents of 4.0 ± 1.1 pA, reached at 2.7 ± 0.7 ms, with 10–90% rise times of 1.4 ± 0.3 ms and decay time constants of 7.7 ± 4.4 ms (range 2.5–16.4 ms; single-exponential fit between 80% and 20% of the peak).

**Figure 1.**
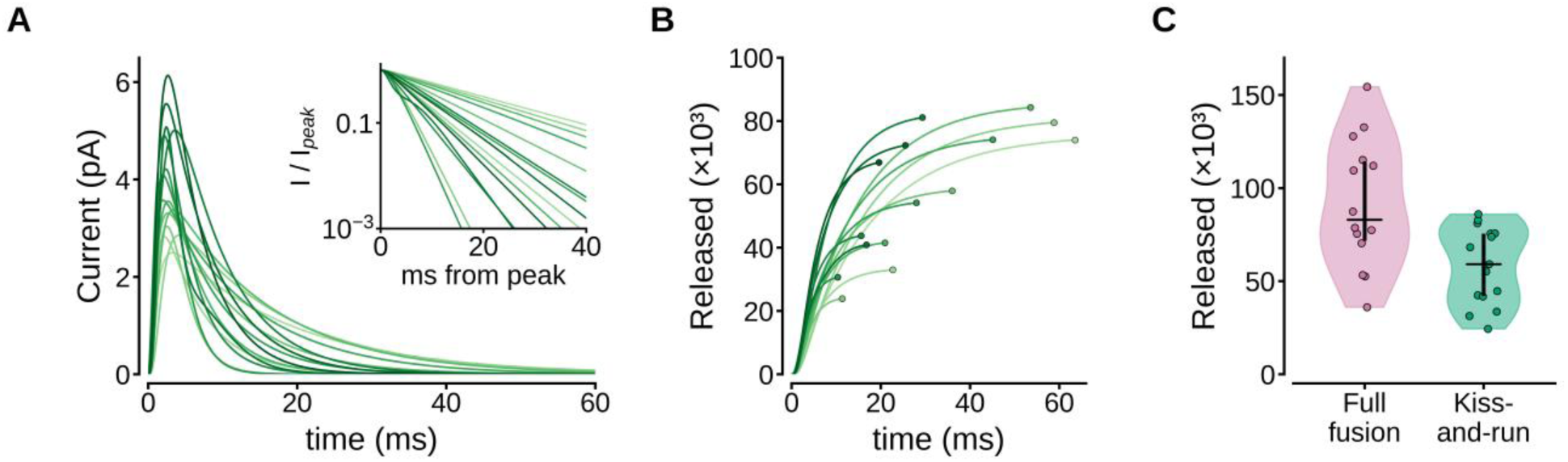
Kiss-and-run events release a fraction of the quantum. (A) Fitted currents of the 15 kiss- and-run events shaded from light (smaller) to dark (larger). Inset, currents normalized to their peaks and plotted on logarithmic scale, showing single-exponential decays. (B) Cumulative number of serotonin molecules released (Equation 10). Traces end upon 98% of release (circle). (C) Serotonin molecules released by full-fusion and kiss-and-run events; violins show kernel density estimates; horizontal bars medians and vertical bars interquartile ranges (Welch’s t-test, p = 0.005).

The cumulative release of each spikelet approached a plateau, reaching 98% of its final value 10–64 ms after onset (Fig. 1 B). Spikelets released 58,344 ± 20,759 molecules (range 24,352–85,961; Shapiro–Wilk *p* = 0.20), as expected if release ceased before the vesicle had completely emptied. By contrast, full-fusion spikes released quanta containing 91,585 ± 34,428 molecules (range 35,964–154,385; *p* = 0.86; Fig. 1 C), a statistically significant difference to Kiss-and-run events (Welch’s *t*-test, *p* = 0.005; Mann–Whitney *U*, *p* = 0.012). On average, spikelets released 64% of the quantum. For leech serotonergic dense-core vesicles 103.7 nm in diameter (Bruns et al., 2000), the full-fusion mean corresponds to an intravesicular serotonin concentration of ∼260 mM, close to the 270 mM estimated by same authors (2000). That these spikelets released only part of the quantum reinforces that they are kiss-and-run events.

### Peak pore diameter estimated by hydrated flux packing (HFP)

Figure 2A shows the optimal packing of 1–10 hydrated serotonin molecules in a circle representing the pore mouth. The enclosing diameter increases in discrete steps with *N*, except between *N* = 6 and *N* = 7, which fit within the same circle. At the current peak, the 15 kiss-and-run events carried 3.9–9.6 molecules/μs, requiring 4–10 hydrated molecules to traverse the pore cross-section simultaneously. The corresponding pore diameters were 2.1–3.2 nm (central values; 1.9–3.4 nm across *d*_H_ = 0.8-0.9 nm; Fig. 2 B; Table 1). These diameters represent lower bounds, because an HFP pore is the narrowest capable of accommodating the peak flux. Packing unhydrated (bare) serotonin molecules (0.67 nm; see Materials and methods) yields diameters 21% smaller (1.6–2.5 nm; Fig. 2 B). The reasons for using the hydrated diameter are given in Materials and methods.

**Figure 2.**
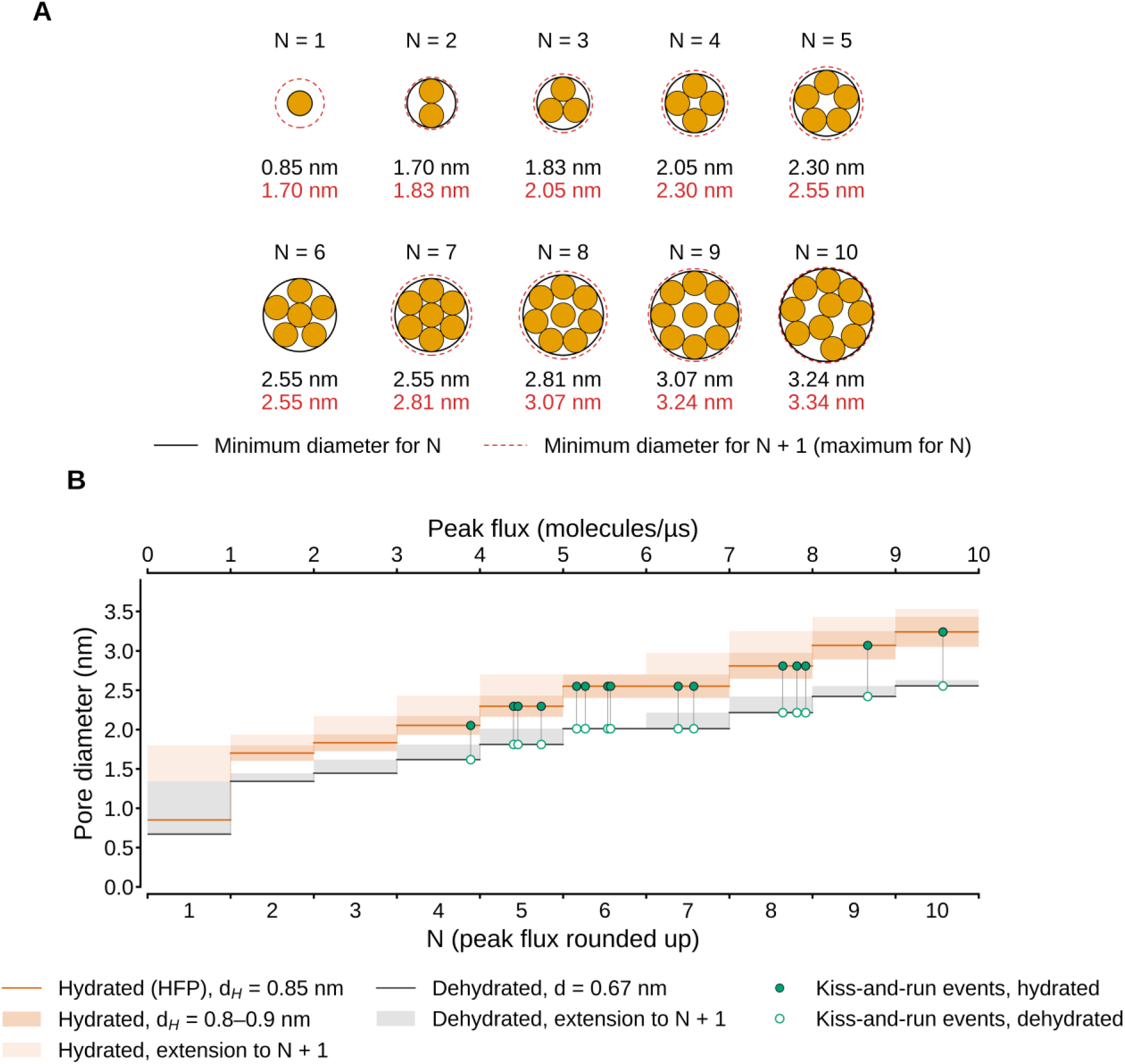
Minimum pore diameter from hydrated flux packing (HFP). (A) Optimal packing of N = 1–10 hydrated serotonin molecules in a circular section (d_H_ = 0.85 nm). Solid black circles, minimum enclosing circle for N molecules (Eq. 11); dashed red circles, minimum enclosing circle for N + 1 molecules, the largest diameter that still corresponds to N. The two diameters are given below each arrangement (black, N; red, N + 1). For N = 6 the two circles coincide. (B) Pore diameter for each value of N (bottom axis), obtained by rounding the peak flux (top axis) up to the next integer, for hydrated serotonin (HFP; orange) and bare serotonin (black; d = 0.67 nm, van der Walls diameter of a sphere). Solid lines, minimum diameter for N; dark orange bands, range for d_H_ = 0.8–0.9 nm; light orange and gray bands, extension to the minimum diameter for N + 1 (d_H_ = 0.9 nm for hydrated molecules). Filled green circles, HFP diameters at peak flux; open circles, same events with bare molecules.

**Table 1.**
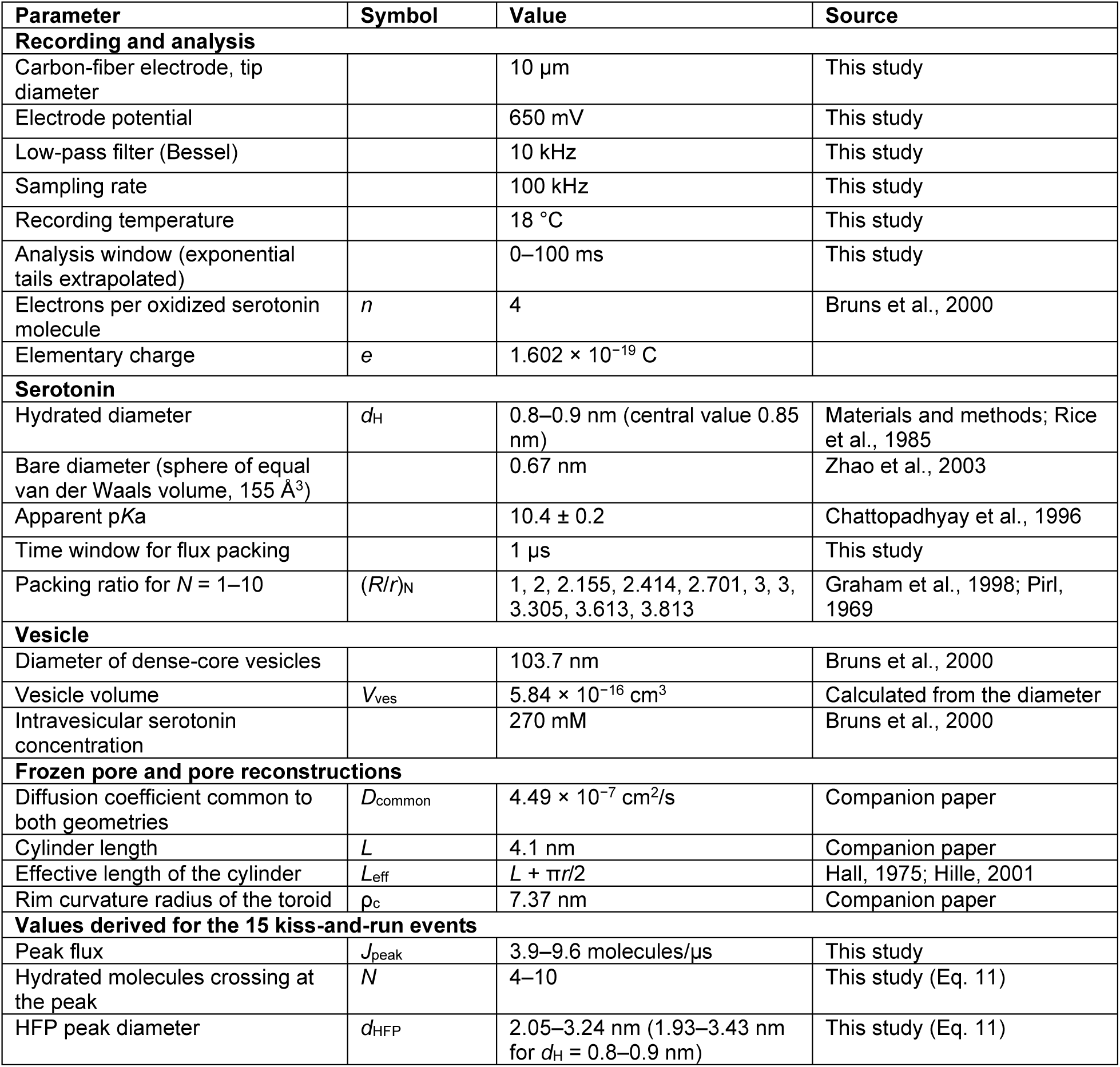
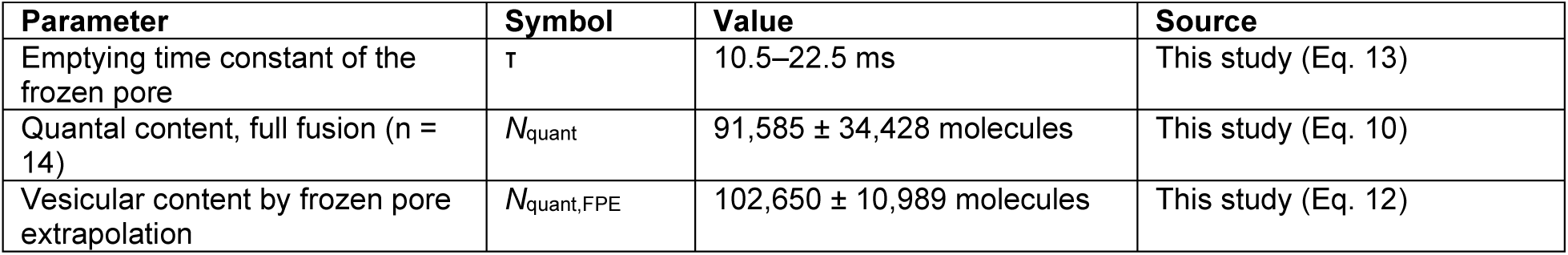
Parameters used in this study. Values labeled “This study” were measured or derived here; ranges of derived values span the 15 kiss-and-run events. Values from the companion paper are those obtained there with the same events.

### Serotonin efflux with fixed pore

With the pore diameter at the current peak in hand, we examined how flux through a pore of constant diameter depends on the quantities that determine it (Eq. 13; Fig. 3). Release through such a pore decays exponentially according to:

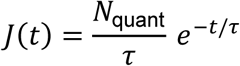

with *τ* = *V*_ves_*L*_eff_/(*Dπr*^2^). The integral of the flux from *t* = 0 to ∞ equals the vesicular quantum *N*_quant_, and the initial amplitude, *N*_quant_*Dπr*^2^/(*V*_ves_*L*_eff_), is proportional to the content and *D* value, and inversely proportional to the vesicle volume and to *L*_eff_/(*πr*^2^). For a pore 2.55 nm wide (median event in Figure 2), *τ* = 15.6 ms, and a vesicle containing 102,650 molecules (the FPE quantum) produces a peak flux of 6.6 molecules/µs. Doubling the content doubles the amplitude without changing *τ* (Fig. 3A). A wider pore increases the initial flux amplitude and shortens *τ* (4.5–9.8 molecules/µs and 22.5–10.5 ms for diameters of 2.05–3.24 nm; Fig. 3B); doubling *D* doubles the amplitude and halves *τ* (Fig. 3C); a larger vesicle with the same content lowers the amplitude and lengthens *τ* (10.7, 6.6, and 4.3 molecules/µs for vesicles 88.1, 103.7, and 120 nm in diameter, respectively; Fig. 3D).

**Figure 3.**
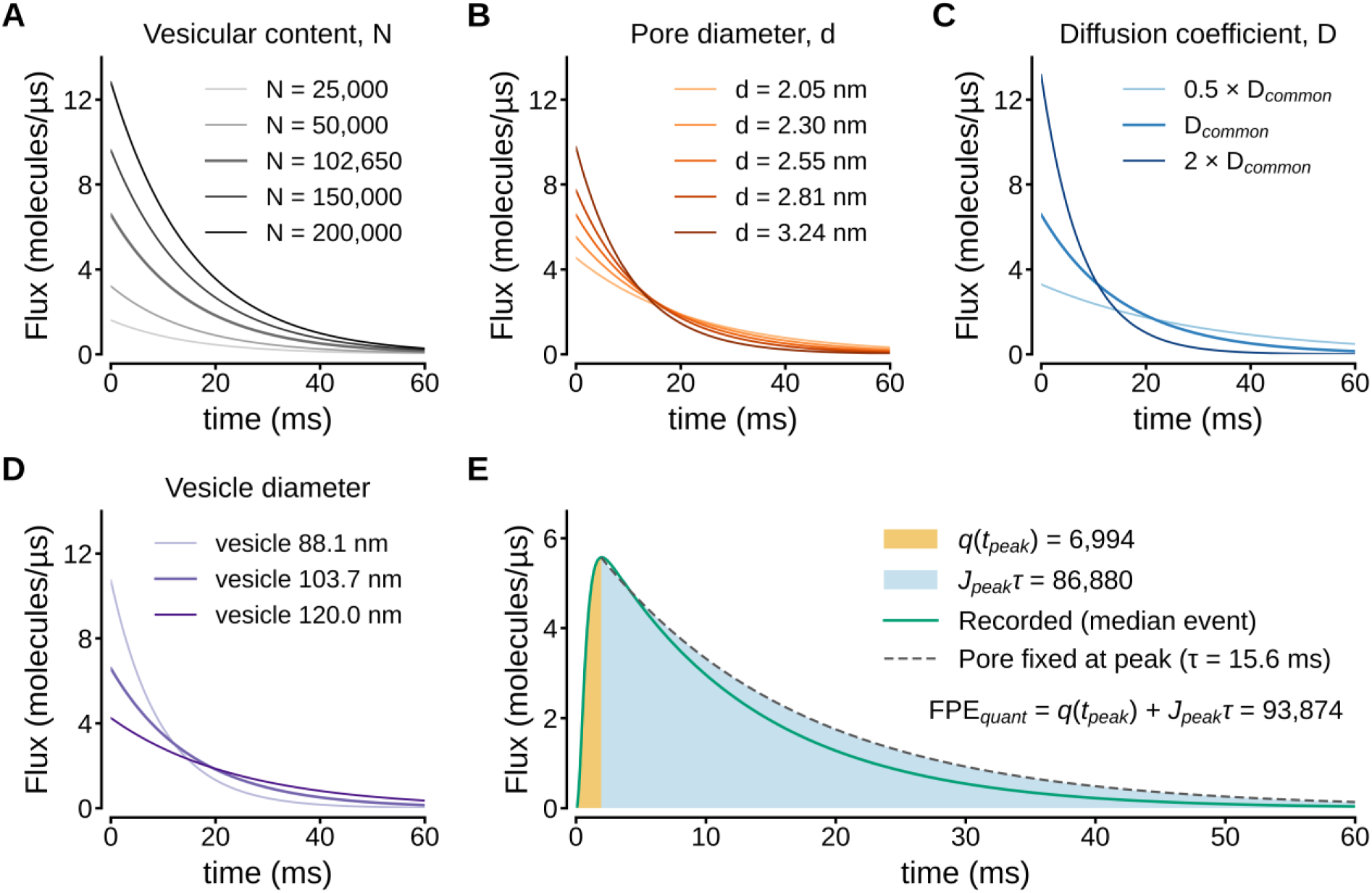
Determinants of serotonin flux through frozen pores. (A) Vesicular content: the amplitude is proportional to N; τ is unchanged; the area under each curve equals N. (B) Wider pores raise amplitude and shorten τ. (C) D_common_. (D) Vesicle diameter. 88.1 and 103.7 nm are axonal and somatic dense-core vesicles of Retzius neurons (Bruns et al., 2000). Larger vesicle lowers the amplitude and lengthens τ. (E) Frozen pore extrapolation (FPE) of the vesicular content of the median kiss-and-run event (Eq. 12). *J*_peak_τ (blue area under the dashed exponential). Green, recorded flux.

Since the vesicular content influences flux through the product *J*_peak_*τ*, it can be recovered from the peak flux if *τ* is known. Eq. 12 adds to the molecules released prior to the peak those that a pore fixed at its peak diameter would release subsequently (Fig. 3E). For the median event, FPE yields 6,994 + 86,880 = 93,874 molecules. The specific pore geometry enters this relationship through *τ*. Using *D*_common_ (companion paper), a toroidal pore (rim curvature radius 7.37 nm) and a short cylinder yield the same *τ* at 2.55 nm. Values were maintained with a small 2.4% variation across the entire range of event diameters (2.05–3.24 nm). Neither the frozen pore fluxes nor the content estimated from them depend on whether the pore geometry is modeled as toroidal or cylindrical.

### The pore closes faster than the vesicle empties

If pores remained at peak diameter, flux decayed exponentially with the time constant of that pore flux (10.5–22.5 ms; Table 1). However, the recorded flux fell faster (Fig. 4, A– C, left axes), reaching half of its peak in 2.3–12.2 ms, compared with 7.3–15.6 ms for frozen pores, representing a ratio of 0.59 in the median event (range 0.23–0.95; two-sided exact Wilcoxon signed-rank test, *n* = 15, *p* = 6.1 × 10^−5^; Fig. 5 A). At one-quarter of the peak the ratio was 0.52.

**Figure 4.**
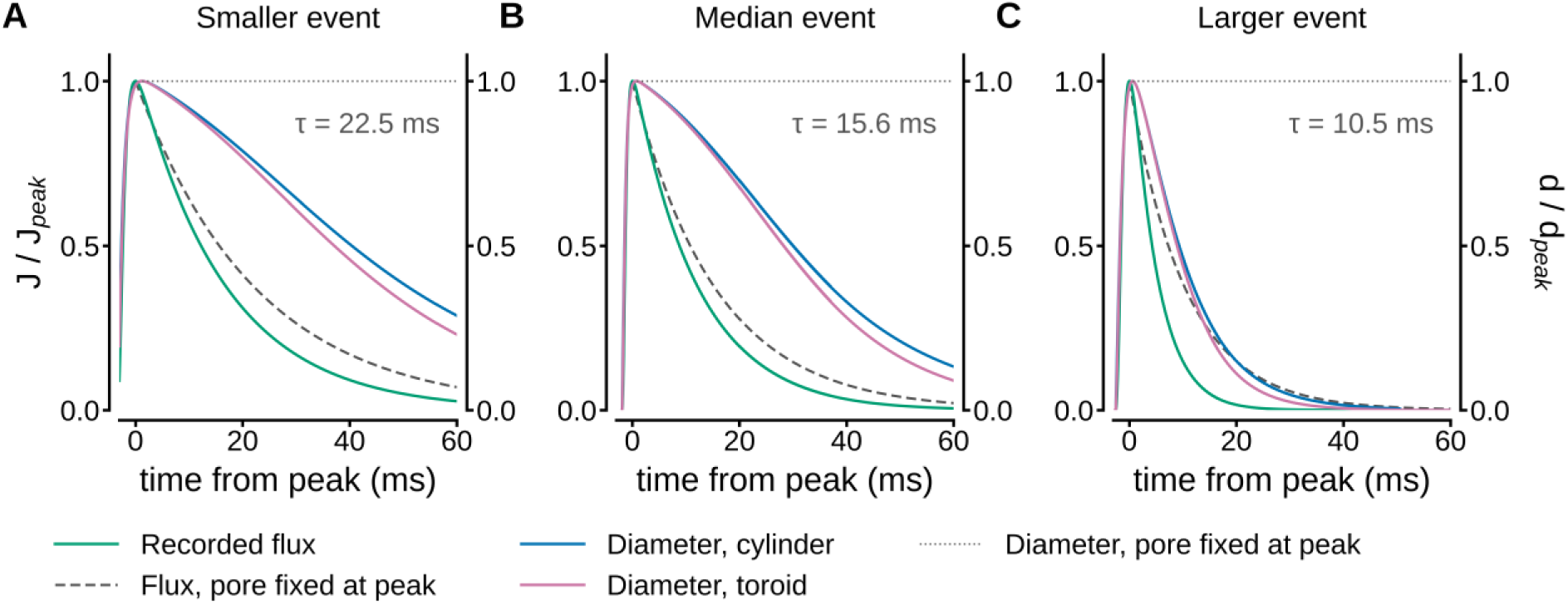
Serotonin flux decays faster than predicted by frozen pore. Left axes: Normalized serotonin recorded flux (green) and flux through a pore frozen at peak HFP diameter (dashed); Right axes: Normalized pore diameter for cylindrical (L = 4.1 nm; blue) and a toroidal (rim curvature radius 7.37 nm; magenta) pores using a common D value (see companion paper. In the larger event the pore closed before flux through frozen pore ends.

**Figure 5.**
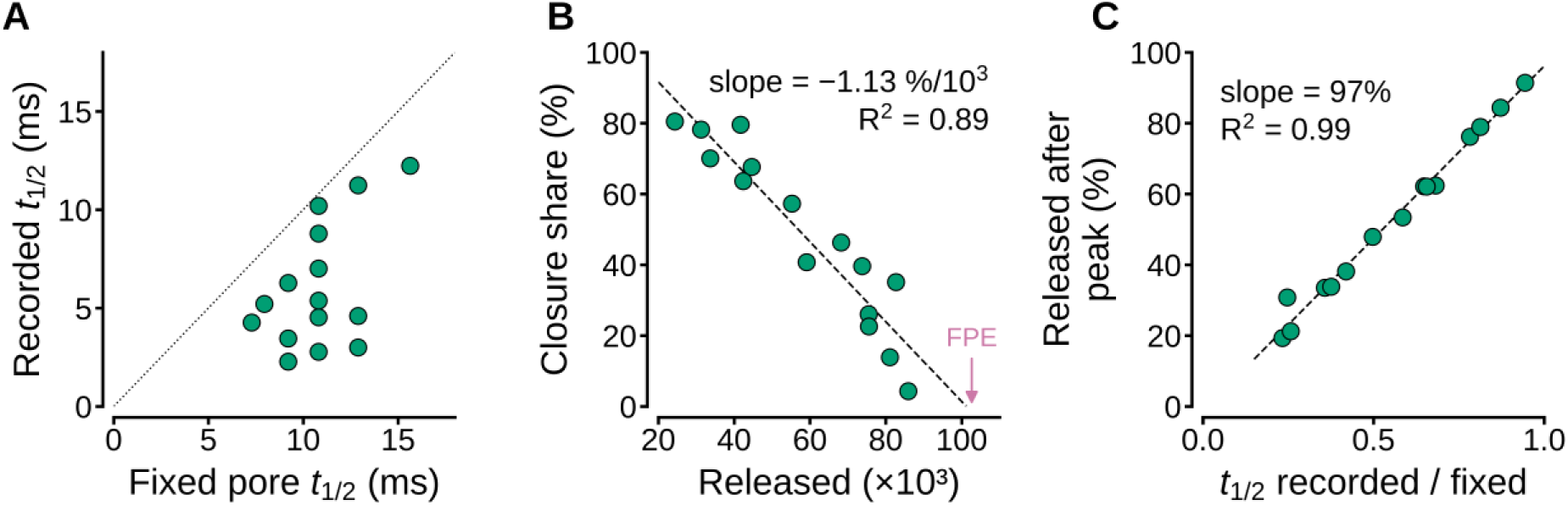
Closure limits kiss-and-run release. (A) Time to half peak for recorded and fixed-pore fluxes of the 15 events; dotted line, identity. (B) Share of flux decrease at half-decay attributable to pore closure versus molecules released. (C) Fraction of remaining content released after peak versus half-decay time ratio of recorded to FOE. B and C use the FPE contents (mean 102,650 molecules).

The faster half decay of the recorded flux implies that the pore narrowed during serotonin release. Reconstructing the pore diameter from the recorded flux evaluated with *D*_common_ and the FPE content of each event makes this behavior explicit (Fig. 4, right axes). The reconstructed diameter reached its maximum 0.6–1.2 ms after the current peak and subsequently decreased. By contrast, a fixed pore would have maintained its peak diameter. When both quantities are normalized to their respective peak values, the diameter decayed more slowly than the flux, reaching half peak values 39.2, 29.2 and 8.7 ms after the diameter peak in the three standard events for a cylinder, and 35.9, 27.4 and 8.3 ms for a toroid. The recorded flux reached half-peak values at 12.2, 8.8, and 4.3 ms after the current peak (see companion paper). This difference arises because the flux falls both as the pore closes and as the vesicle empties (Fig. 5; Eq. 14). This means that a modest narrowing of the pore causes the recorded flux to fall faster than that of a fixed pore. Both geometries close along nearly identical time courses, the toroid closing slightly faster. This demonstrated active pore closure independently of its geometry.

When the decay of the flux was separated into depletion and pore-resistance terms (Eq. 14), using the FPE content of each event (Materials and methods), the increase in pore resistance accounted for 46% of the drop in flux at half-decay (median event; range 4– 81%). Events in which pore closure dominated released the smallest amount of serotonin (Spearman *ρ* = −0.96, p = 1.3 × 10^−8^; Fig. 5 B). The fraction of the remaining content released after the peak (19–91%) increased with the ratio between the recorded and fixed-pore half-decay times (*ρ* = 0.99; Fig. 5C). Thus, the earlier the pore closed relative to the emptying time constant of the vesicle, the smaller the total fraction of serotonin released.

### Kiss-and-run vesicles contain a full serotonin quantum

Frozen pore extrapolation (Eq. 12) estimates the serotonin content of each kiss-and-run vesicle as the molecules released before the flux peak, *q*(*t*_peak_), plus those that the pore, frozen at its HFP diameter, would release afterwards, *J*_peak_τ (Fig. 3 E). *q*(*t*_peak_) contributed 6,000–18,000 molecules and *J*_peak_τ 80,500–105,100 molecules (Fig. 6 D). The FPE contents averaged 102,650 ± 10,989 molecules (range 88,110–121,707), similar (p = 0.27; 95% confidence interval of the difference was −9,405 to +31,535) to the mean quantum released by full-fusion events (91,585 molecules; Welch’s t-test, p = 0.27; Fig. 6, A and B). Vesicles undergoing kiss-and-run were therefore fully loaded, and their smaller release did not reflect a smaller quantum.

**Figure 6.**
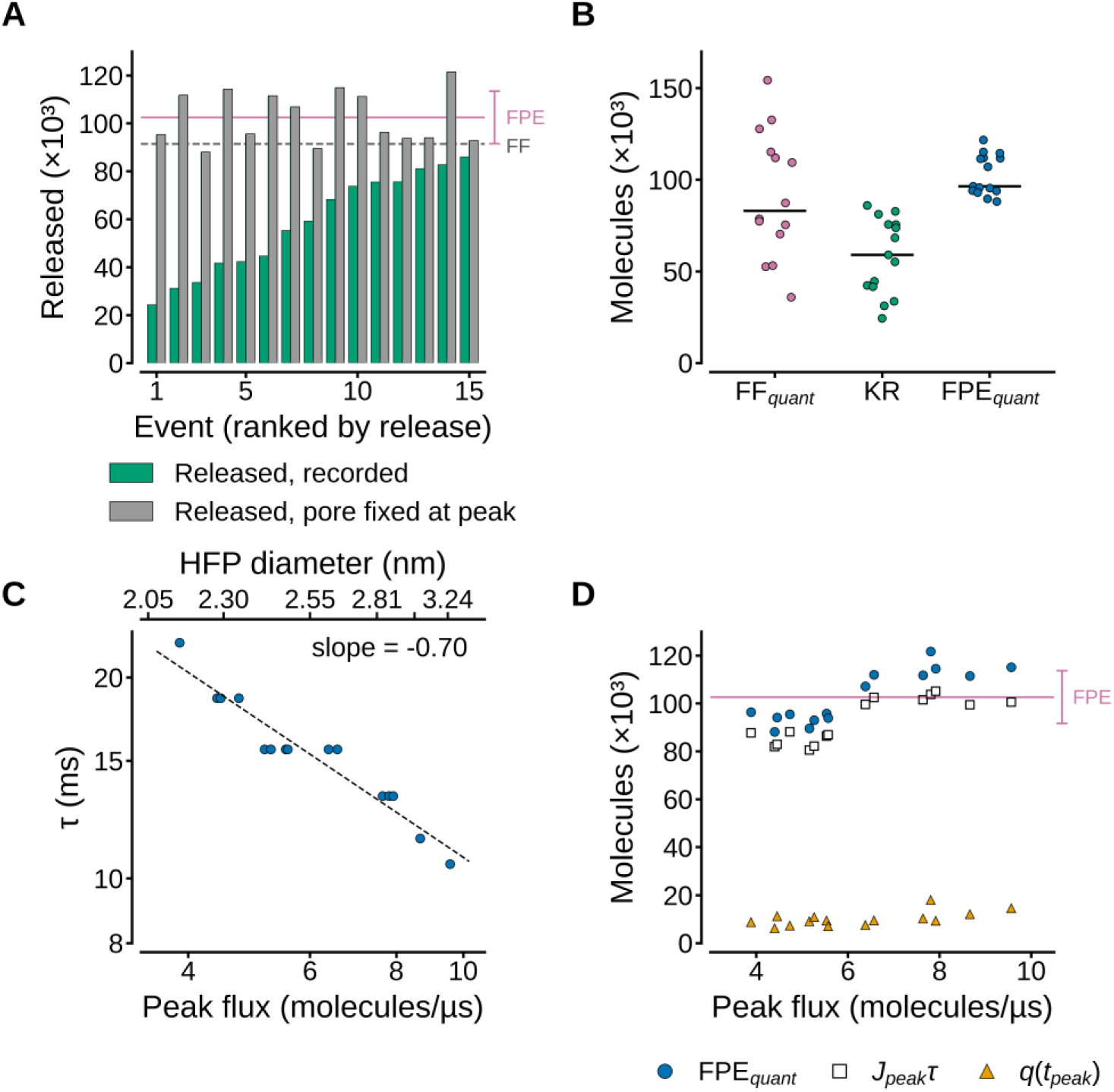
Vesicular content estimated by FPE. (A) Serotonin molecules released (green) and FPE content of the same event (gray); solid line and error bar, mean ± SD of the FPE quantum; dashed line, mean quantal size of full-fusion (FF) events. (B) Full-fusion quantum (FF_quant_), molecules released by kiss- and-run (KR), and FPE quantum (FPE_quant_; Eq. 12); horizontal lines, medians. (C) Emptying time constant of pores of HFP diameter versus peak flux (log–log); dashed line, power-law fit; upper axis, HFP diameter. (D) FPE content (circles), *J*_peak_*τ* (squares) and molecules released before peak (triangles) versus peak flux for every event; solid line and error bar, mean ± SD of the FPE quantum.

The FPE contents varied less than the full-fusion integrals (coefficients of variation, 11% and 38%; Fig. 6 B). Because the HFP diameter increased with peak flux, the time constant of the frozen pore decreased as peak flux increased (τ α *J*_peak_^−0.70^; Fig. 6 C). *J*_peak_τ varied ∼10% between events, compared with 28% for the peak flux. The estimate therefore establishes the content of the population, whose scale is set by the D value more reliably than that of each vesicle (see Discussion). Kiss-and-run events released 26–92% of their vesicular serotonin relative to their FPE content (Fig. 6 A). The events that released the smallest fractions had faster flux relative to that of the frozen pore (Fig. 5 B).

## Discussion

Our results show that kiss-and-run serotonergic events in Retzius neurons occur from fully loaded vesicles in which the pore closes, preventing transmitter depletion. About two-thirds of the quantum is released through pores that expand and close.

To our knowledge, this is the first study that determines for the same kiss-and-run events, the size of the pore, the quantal content of the vesicle and the process that limits release. The hypothesis that kiss-and-run releases only part of the quantum is well established (Alvarez de Toledo et al., 1993; Staal et al., 2004; MacDonald et al., 2006; Alabi and Tsien, 2013). Electrochemical cytometry has shown that, on average, vesicles release about 40% of their content (Omiatek et al., 2010). These fractions however compare populations of isolated vesicles with populations of exocytotic events. Therefore, they cannot determine whether individual subquantal events arise from a partially filled vesicle, from a pore that remains open while the vesicle empties, or from a pore that closes early. Here, sizing the pore with HFP, estimating the content of each vesicle by frozen pore extrapolation and comparing each event with its frozen pore separated those possibilities. Vesicles were fully loaded and closure of the pore set the amount released.

### Hydrated flux as external standard for pore diameter

HFP provides a lower bound for the pore diameter that depends solely on the measured flux and the size of hydrated serotonin. It does not require the parameters that render conductance, diffusion coefficient, pore geometry or the vesicular content, or amperometry-based estimates of pore size model-dependent (Breckenridge and Almers, 1987a; Oleinick et al., 2013). The method assumes that the molecules crossing the pore within one microsecond occupy its cross-section simultaneously; this transit time sets the absolute scale. The resulting diameters, 2.1–3.2 nm, fall within the range of non-dilating fusion pores and serve as reference in the companion paper to compare toroidal and cylindrical pore geometries.

### Bare serotonin diameters for HFP

Some reasons for using the hydrated diameter of serotonin were given above. Here it suffices to mention that the bare molecular diameter would lower the HFP bound by 21% (Fig. 2 B), under the assumption of a fully dehydrated transit, which a 2–3 nm lipidic pore lacking coordinating groups does not impose (Hille, 2001). Partial dehydration might occur transiently while pores are narrower than ∼1 nm, either during opening or closing. By contrast, the HFP procedure is evaluated exclusively at the flux peak.

### Quantal content estimate

The quantal values measured here for full-fusion (91,585 molecules) and FPE contents (102,650 molecules), match the ∼ 95,000 molecule content expected for a 103.7-nm vesicle filled with 270 mM serotonin (Bruns et al., 2000). Factors that account for the upper level variability are that our full-fusion events were selected as spikes exceeding five times the baseline noise, thus excluding smaller events within the broad charge distributions reported by those authors (2000). Moreover, currents here were integrated after extrapolating the exponential tails to 100 ms. This approximation adds the late-decay charge that is otherwise lost to baseline noise. Finally, the mean of 14 events drawn from a distribution with a coefficient of variation of 38% carries a standard error of about 10%.

### Serotonin flux versus pore closure

Comparing serotonin measured flux to FPE flux as indicators of pore closure is independent of vesicular content, because the emptying time constant of a frozen pore depends exclusively on its geometry and D value. The faster decay in every recorded current demonstrates that the pore actively narrowed while serotonin release was ongoing. This finding aligns with the accelerated closure of toroidal pores described in our companion paper and with reconstituted fusion pores that flicker and close without undergoing full dilation (Bao et al., 2018). While time courses are smoothed by diffusion between the pore exit and the electrode (Zhang and Jackson, 2008), affecting both recorded and fixed-pore currents similarly near the peak, the spatial diffusion might slightly prolong the decay tail in our recordings; if so, the speed of closure is underestimated.

## Data availability

The amperometric events analyzed here, the fitted currents, the parameter values and the Python code used for the calculations and figures are available from the corresponding author upon reasonable request.

## Acknowledgments

The kiss and run current traces in this and in the companion paper were extracted from amperometric recordings made for other purposes by Mrs. Montserrat Cercós in my laboratory. I thank Sara Flores González for her excellent assistance with animal and laboratory care and Francisco Pérez from the Computer unit of IFC-UNAM, for technical support. This study was funded by a grant from the Human Frontier Science Program Organization (RGP0060/2019–102) to F.F. De-Miguel and by grant AG200823 from the Dirección General de Asuntos del Personal Académico, UNAM, to F.F. De-Miguel and Martha Takane.

## Abbreviations

5-HT: 5-hydroxytryptamine (serotonin)
FF: full fusion
HFP: hydrated flux packing
KR: kiss-and-run
SEMD: statistical empirical mode decomposition.

## Supplemental material

### Supplemental text. Release through a pore of constant diameter

This text derives the properties of the fixed-pore model used in Eqs. 12 and 13 and in Figs. 3 and 5, and verifies them numerically with the parameters of this study: a cylindrical pore 2.55 nm in diameter (the HFP diameter of the median event) and 4.1 nm long, *D* = *D*_common_ = 4.49 × 10^−7^ cm²/s, and a vesicle 103.7 nm in diameter (*V*_ves_ = 5.84 × 10^−16^ cm³).

**S1.** *Governing equation.* A pore of constant geometry connects a well-mixed vesicle containing *N*(*t*) molecules to a perfect extracellular sink. By Fick’s law, the flux through the pore is the concentration difference, *N*(*t*)/*V*_ves_, divided by the diffusional resistance of the pore, *R* = *L*_eff_/(*D*π*r*^2^), where *L*_eff_ = *L* + π*r*/2 includes the access resistance of the two mouths (Hall, 1975; Hille, 2001). The flux is therefore proportional to the content,

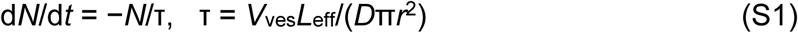

with the solution *N*(*t*) = *N*_0_exp(−*t*/τ), where *N*_0_ is the content when the pore reaches its constant size, and the current is *I*(*t*) = (*neN*_0_/τ)exp(−*t*/τ). The time enters only through *D*, whose units (cm²/s) carry the rate of Brownian motion; the gradient that drives the flux is spatial, across the length of the pore.

**S2.** *The time constant is a property of the pore.* τ contains only the vesicle volume, the geometry of the pore and the diffusion coefficient, and not the content. For the pore defined above, τ = 15.6 ms. Fitting ln *I*(*t*) with a straight line on currents computed for *N*_0_ = 25,000 to 200,000 molecules recovered τ = 15.597 ms in every case, identical to the value given by the formula (Table S1). A larger content does not make molecules cross faster: it increases the number of molecules near the pore at any instant, and with it the flux, but not the fraction of the content that leaves per unit time.

**S3.** *The amplitude is proportional to the content.* With the pore fixed, the initial current, *neN*_0_/τ, is proportional to *N*_0_: 1.03, 2.05, 4.22, 6.16 and 8.22 pA for 25,000, 50,000, 102,650, 150,000 and 200,000 molecules (Table S1; Fig. 3 A).

**S4.** *Mass conservation.* The integral of the current, divided by *ne*, equals *N*_0_ exactly over infinite time. Over the 100-ms window used in this study it recovers 99.8% of the content for all values of *N*_0_ (Table S1), the remainder lying in the tail beyond 100 ms. A larger content produces a proportionally taller current and a proportionally larger total release, so that exactly *N*_0_ molecules leave.

**Table S1.** Current through a fixed pore for different vesicular contents. Pore 2.55 nm in diameter and 4.1 nm long, *D*_common_, vesicle 103.7 nm in diameter; τ from the formula, 15.597 ms. Time to 100 molecules, τ ln(*N*_0_/100).

| **N<sub>0</sub> (molecules)** | **Initial current (pA)** | **τ fitted (ms)** | **Released in 0–100 ms (molecules)** | **Time to 100 molecules (ms)** |
|---|---|---|---|---|
| 25,000 | 1.03 | 15.597 | 24,959 (99.84%) | 86.1 |
| 50,000 | 2.05 | 15.597 | 49,918 (99.84%) | 96.9 |
| 102,650 | 4.22 | 15.597 | 102,481 (99.84%) | 108.2 |
| 150,000 | 6.16 | 15.597 | 149,754 (99.84%) | 114.1 |
| 200,000 | 8.22 | 15.597 | 199,671 (99.84%) | 118.6 |

**S5.** *Two meanings of decay time.* The time constant τ, the time for the current to fall by a factor *e* from any instant, does not depend on the content. The time needed to drain the vesicle to a fixed absolute number of molecules does, but only logarithmically: *t* = τ ln(*N*_0_/*N*_threshold_). For the median event, the 86,880 molecules remaining at the current peak would fall to 100 molecules in 105.5 ms; a content 50% larger would take 111.9 ms, 6% longer. This distinction is the reason why the comparison of recorded and fixed-pore currents through their times to fall to one-half of the peak (*t*_1/2_ = τ ln 2 for the fixed pore; Fig. 5 A) is independent of the vesicular content assumed for each event.

**S6.** *Estimation of the content (*Eq. 12). Because τ does not depend on the content, a fixed pore that reproduces the measured peak current must contain *I*_peak_τ/*ne* molecules at the peak; adding the molecules released before the peak, *q*(*t*_peak_), gives Eq. 12. The estimate depends on the content only through the measured current and on the pore only through τ, so that any error in *D* or in the geometry of the pore scales the estimate of all events by the same factor.

**S7.** *Pore geometry.* All the properties above follow from the first-order kinetics of S1 and do not depend on the expression used for *L*_eff_. With *D*_common_, a toroidal pore with rim curvature radius 7.37 nm gives the same τ as the 4.1-nm cylinder at 2.55 nm and values within 2.4% over the HFP diameters of the 15 events (2.05–3.24 nm; τ = 10.5–22.5 ms), so that neither the fixed-pore currents nor the content estimated from them depend on whether the pore is toroidal or cylindrical.

